# Global cognitive performance is not influenced by diurnal rhythm

**DOI:** 10.1101/2020.12.29.424677

**Authors:** Celine H. De Jager, Eleanna Varangis, Yaakov Stern

## Abstract

**Objective:** The human brain is influenced by different biological rhythms, including the daily 24-hour (diurnal) cycle. Past studies have reported evidence of variation in cognitive performance over the course of the day, and of differences in the peak time for cognition in older age. Here, we investigated these questions using two existing longitudinal studies of healthy adults.

**Methods:** Time of neuropsychological battery testing was extracted from study records, and we analyzed cognitive performance measures from 4 domains (Vocabulary, Processing Speed, Fluid Reasoning, and Episodic Memory) in 543 healthy adults between the ages of 20 and 80. Time of day was dichotomized as morning (281 tested before noon), and afternoon (242 tested after noon).

**Results:** Multivariate analyses controlling for both gender and years of education revealed no significant effect of time of testing (or its interaction with participant age) on cognitive performance. These results suggest that diurnal effects during time periods typically used to test human subjects are unlikely to have a meaningful effect on performance on the neuropsychological tests that are used for standard cognitive assessment.

**Conclusion:** This suggests that the effect of time of day on cognition in the context of aging may not be as ubiquitous as previously suggested, and thus is unlikely to represent a large confound in existing studies of cognition across the adult lifespan.

## Introduction

Cognitive performance is carried out by the brain, which, like the rest of the body, is influenced by different biological rhythms, including the daily 24-hour cycle. This rhythm is linked to our sleep cycle and is driven by our internal circadian clock. This clock is best described as a biological pacemaker with a period of about one day, and several different hormones carry out this cycle under the direction of the internal clock. Intuitively, we may feel that our own cognitive performance varies over the course of the day, and such an effect could have profound effects on the interpretation of results from studies that do not take into account such an effect. Further, we may wish to select specific times of the day for testing in order to ensure that participants are performing at their optimal level.

Some studies have reported evidence that circadian effects may vary with age: older adults’ peak time of functioning may be in the morning, while younger adults’ peak time of functioning may be later in the day (Adan & Almirall, 1990; Bodenhausen, 1990; Colquhoun, 1971; Folkard, Knauth, Monk, & Rutenfranz, 1976; Folkard, Weaver, & Wildgruber, 1983; Mecacci, Zani, Rocchetti, & Lucioli,1986; Wilson, 1990). This is better known as the “synchrony effect.” Thus, it may be important to account for this shift in biological rhythm that appears to begin at around age 50 (Ishihara, Miyake, Miyasita, & Miyata, 1991). Other investigators have also found, using a Morningness-Eveningness Questionnaire, that there are significantly higher morningness ratings for older adults than for younger adults (Hoch et al., 1992; Mecacci, Zani, Rochetti, & Lucioli, 1986). Synchrony between optimal performance periods and the times at which testing is conducted may be a critical variable when assessing group differences in performance, especially when examining differences in cognitive performance between older and younger adults (May, Hasher, & Stoltzfus, 1993).

Many investigators (such as May et al., 1993) have explored different types of cognitive processes that may be affected by one’s peak circadian arousal period and the time at which testing occurs. These reports suggest that circadian arousal is correlated with performance on a variety of tasks, and that performance peaks at a certain level of circadian arousal. Further, in one study, dramatic differences in memory performance were noted across times of day in both younger and older adults, and these patterns of performance throughout the day were quite different for younger and older adults (May, Hasher, & Stoltzfus,1993). More recently, older adults tested in the morning were reported to be better able to ignore distractions than those tested in the afternoon (Anderson, 2014). However, these reports are balanced by a number of other studies which have found that there had been no effect on time of day on working memory performance (Sunram-Lea, 2001). Additionally, most of the studies that reported effects of synchrony on cognitive performance were performed in small samples.

Thus, the extent to which time of day influences cognition in those periods typically used to test human subjects remains unclear, and it is essential to know whether time of day is an important covariate in study design. Many of the studies mentioned above use attention network-based tasks which may be more vulnerable to circadian effects than other cognitive tasks that are frequently used in human studies, as well as in a clinical setting. Thus, we decided to compare the effect of time of day on performance in three cognitive domains affected by aging (Processing Speed, Fluid Reasoning, and Episodic Memory) as well as in an unaffected domain (Vocabulary) in a sample of nondemented adults between the ages of 20 and 80.

## Methods

### Participants

Subjects were drawn from two long-term observational studies of aging and cognition, the Reference Ability Neural Network (RANN) and Cognitive Reserve (CR) studies (Habeck, C. et al., 2017; Habeck, C. et al., 2012; Stern, Y. 2007). Participants were recruited through random market mailing procedures, flyers, and word of mouth. Interested respondents replied by mail, telephone, or internet. Potential respondents were first screened by telephone, and then invited for an in-person screening. All of the participants were healthy, right-handed, and without mild cognitive impairment (MCI) or dementia. The inclusion criteria were designed to rule out people with contraindications for magnetic resonance imaging (MRI), hearing or vision impairments, medical or psychiatric conditions that could influence performance, or medications that target the central nervous system. Cognitive task data was available for 543 healthy adults between the ages of 20 and 80. Participants also provided information about ethnicity, gender, and years of education.

### Time of Day

The scheduling of the study visits was not randomized; participants were scheduled based on their availability.

Manual review of the study records for each of the 543 subjects was performed, and the time of each study visit was recorded. For analyses, study visits were binned into either morning (before noon) or afternoon (noon and later) visits.

### Cognitive Testing

All participants were tested using a neuropsychological battery designed to measure a wide range of cognitive functions. Selected tasks were administered in the following fixed order: Wechsler Adult Intelligence Scale (WAIS-III; Wechsler, 1997), Letter-Number Sequencing, American National Adult Reading Test (AMNART; Wechsler, 1997), Selective Reminding Task (SRT) immediate recall (Buschke and Fuld, 1974), WAIS-III Matrix Reasoning (Wechsler, 1997), SRT delayed recall and delayed recognition (Buschke and Fuld, 1974), WAIS-III Digit Symbol (Wechsler, 1997), Trail-Making Test versions A and B (TMT-A/B; Reitan, 1978), Controlled Word Association (C-F-L) and Category Fluency (animals; Benton et al., 1983), Stroop Color Word Test (Golden, 1975), Wechsler Test of Adult Reading (WTAR; Holdnack, 2001), WAIS-III Vocabulary (Wechsler, 1997), and WAIS-III Block Design (Wechsler, 1997). Prior analyses using these tasks in our lab demonstrated that these selected tasks reflect four latent constructs (Razlighi et al., 2017): vocabulary (vocab; WAIS Vocabulary, AMNART, WTAR), perceptual speed (speed; WAIS Digit Symbol, 2 measures from the Stroop test, TMT-A), fluid reasoning (fluid; WAIS Matrix Reasoning, WAIS Block Design, and TMT-B), and episodic memory (mem; 3 measures from the SRT).

Following collection of all baseline cognitive data, performance on each task was z-scored relative to the mean and standard deviation for each task within the whole sample of participants enrolled in the RANN and CR studies who completed these assessments. The z-scores for the measures within each cognitive domain were then averaged in order to generate domain z-scores.

### Covariates

Education was measured as years of completed education. Gender was self-reported by participants. All analyses of cognitive task performance controlled for years of education and gender. Further, participant age was transformed into a categorical variable in order to more closely mimic previous studies that looked specifically at differences between older and younger samples. In the first analysis, age was trichotomized to represent younger (age 20-39), middle-aged (age 40-60), and older (age 61-80) age groups in order to probe effects seen when comparing older and younger adults. In the second analysis, age was dichotomized to represent younger (age 20-49) and older (age 50-80) adults, since there is evidence to suggest that circadian rhythms begin to shift around 50 years of age (Ishihara, Miyake, Miyasita, & Miyata, 1991).

### Statistical Analyses

Two multivariate analyses of covariance (MANCOVA) tested for differences in cognition based time of day and age. In each analysis the within-subjects factor (or repeated measures factor) was task domain (vocabulary, processing speed, fluid reasoning, episodic memory), and the between-subjects factors were time of day (before 12pm vs. after 12pm), and participant age (with two different stratifications, as mentioned above and detailed below). The primary outcome measures were the effect of time of day on cognitive task performance, and the effects of interest included the interactions among the three aforementioned factors (Age*Domain, Age*Time, Domain*Time, Age*Domain*Time). Analyses were conducted using SPSS Statistics.

#### Model #1

This MANCOVA evaluated the effects of time of day (AM vs. PM), age (20-39, 40-60, and 61+), and task domain (vocab, speed, fluid, mem - within subjects) on cognitive task performance. The analysis included years of education and gender as silent covariates.

#### Model #2

This MANCOVA evaluated the effects of time of day (AM vs. PM), age (under 50 vs. over 50), and task domain (vocab, speed, fluid, mem - within subjects) on cognitive task performance. This model also included years of education and gender as silent covariates.

## Results

The demographic characteristics of our subjects are presented in **Table 1**. The distribution of the time of neuropsychological testing is bimodal. Thus, we elected to study time of administration as a binary trait. By splitting participants into those tested before or after 12:00, we see that there were roughly equal sample sizes of participants tested in the morning (n=281) as in the afternoon (n=242), ensuring reasonable statistical power for our comparison.

**Table 1:** Subject Characteristics

|  |  |
| --- | --- |
| Gender |  |
| Males (n) | 238 |
| Females (n) | 305 |
| Average Age(years) | 55 (20-80) |
| Average Years of Education (range) | 16 (9-24) |
| Appointment Times |  |
| AM (8:00AM-11:59AM) | 281 |
| PM (12:00PM-5:00PM) | 242 |

In Model #1, the 4 (Domain: Vocabulary, Processing Speed, Fluid Reasoning, Episodic Memory) x 3 (Age Group: 20-39 years old, 40-60 years old, 61-80 years old) x 2 (AM: morning vs. afternoon testing) MANCOVA (covariates: years of education, gender) revealed significant main effects of Domain (F_3,1545_=27.963, p<.001) and Age Group (F_2,515_=60.492, p<.001), as well as significant interactions between Domain and Age Group (F_6,1545_=62.944, p<.001) and Domain and AM (F_3,1545_=3.294, p=0.020)(Figure 2). The main effect of AM (F_1,515_=0.001, p=0.970), the interaction between Age Group and AM (F_2,515_=0.894, p=0.410), and the 3-way interaction among Domain, Age Group, and AM (F_6,1545_=1.353, p=0.230) were not significant, as seen in Tables 1 and 2. The main effect of Domain revealed that, overall, participants performed worse on Vocab tasks than all other tasks (SPEED mean difference=-0.198, p<.001; FLUID mean difference=-0.183, p<.001; MEM mean difference=-0.179, p=0.002). The main effect of Age Group revealed that all three groups differed, such that Younger adults performed better on all tasks than Middle-Aged adults (mean difference = 0.337, p<.001) and Older adults (mean difference=0.586, p<.001), and Middle-Aged adults performed better on all tasks than Older adults (mean difference=0.248, p<.001). The interaction between Domain and Age Group showed that older adults performed better on Vocab tasks than younger and middle-aged adults (F_2,525_=17.961, p<.001), but for all other tasks, younger adults performed better than middle-aged adults, who performed better than older adults (Speed: F_2,525_=104.869, p<.001; Fluid: F_2,526_=61.208, p<.001; Memory: F_2,522_=55.123, p<.001; seen in Table 4 and Figure 3).These provide a good positive control for our analyses. However, the primary goal of our analysis was to evaluate the time of day on cognitive performance, and we find that there is no main effect of time of day (F1,515=0.001, p=0.970). Further, the interaction between Age Group and time of day (F2,515=0.894, p=0.410), and the 3-way interaction among Domain, Age Group, and time of day (F6,1545=1.353, p=0.230) were not significant, as seen in Tables 2 and 3. Thus, while we report significant differences in cognitive performance between subjects of different age groups, the time of day during which testing occurs does not introduce a significant variation in participant performance in Model #1.

**Table 2:**
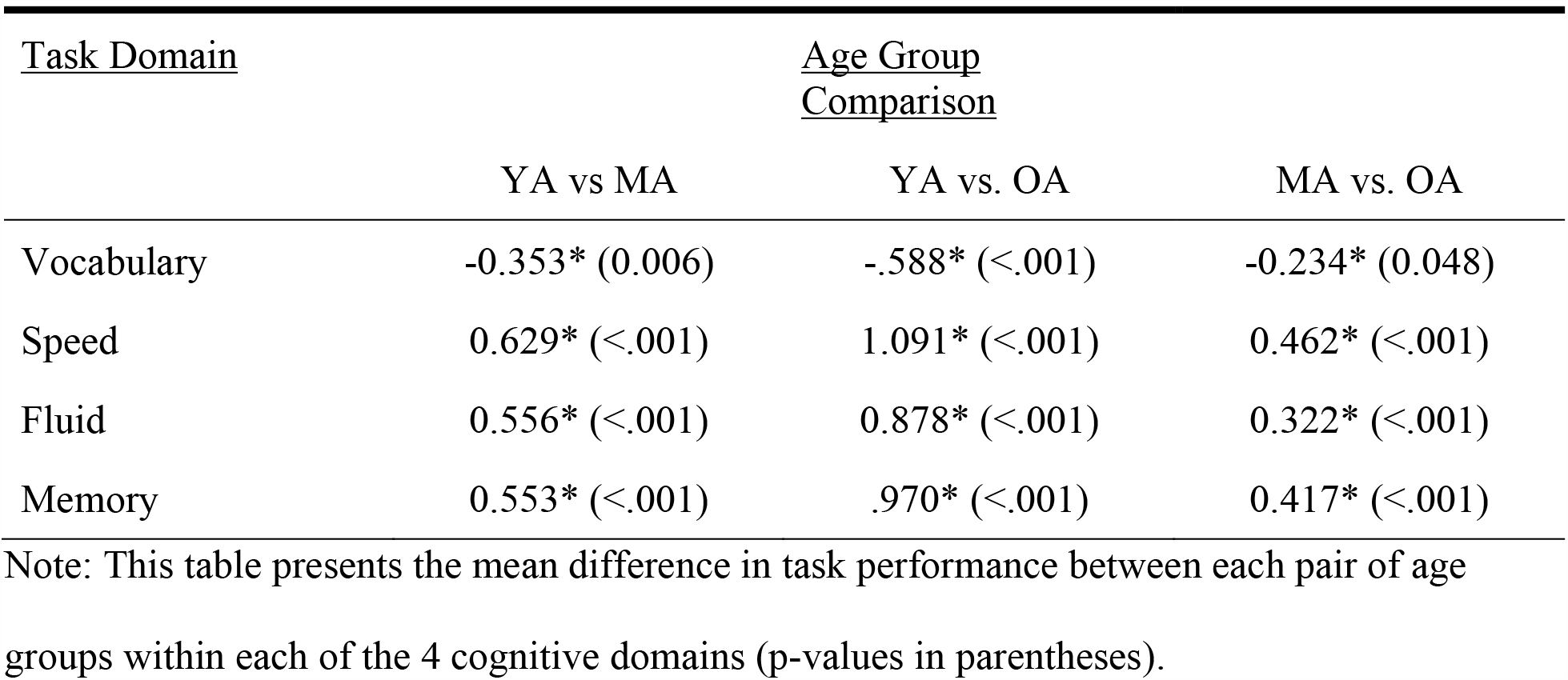
Interaction between Domain and Age Group for Model #1

| <u>Task Domain</u> | <u>Age Group Comparison</u> |  |  |
| --- | --- | --- | --- |
|  | YA vs MA | YA vs. OA | MA vs. OA |
| Vocabulary | -0.353* (0.006) | -.588* (<.001) | -0.234* (0.048) |
| Speed | 0.629* (<.001) | 1.091* (<.001) | 0.462* (<.001) |
| Fluid | 0.556* (<.001) | 0.878* (<.001) | 0.322* (<.001) |
| Memory | 0.553* (<.001) | .970* (<.001) | 0.417* (<.001) |

Note: This table presents the mean difference in task performance between each pair of age groups within each of the 4 cognitive domains (p-values in parentheses).

**Figure 1:**
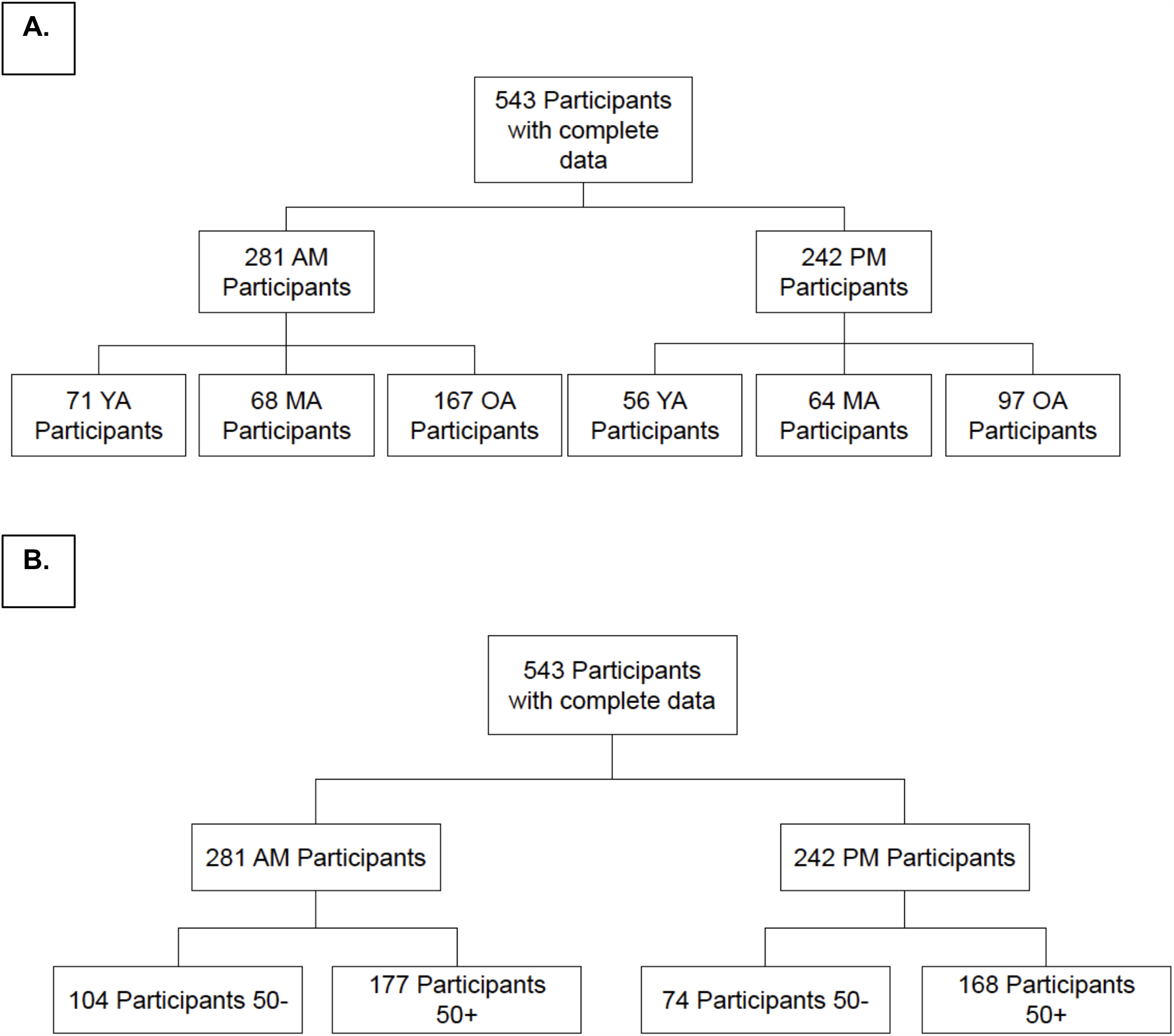
Distribution of participants into the different subgroups in the two analytic models. (A) In Model #1, the 543 participants with complete data were sorted into 6 groups based first on time - before noon (AM) and afternoon (PM) – and then on three age groups - young adults (YA, 20-39), middle aged adults (MA, 40-60), and older adults (OA, 61+). (B) In Model #2, the 543 participants with complete data were sorted into 4 subgroups based first on time - before noon (AM) and afternoon (PM) – and then on two age groups - under the age of 50 (<50) and subjects who were 50 years and older (50+).

**Figure 2:**
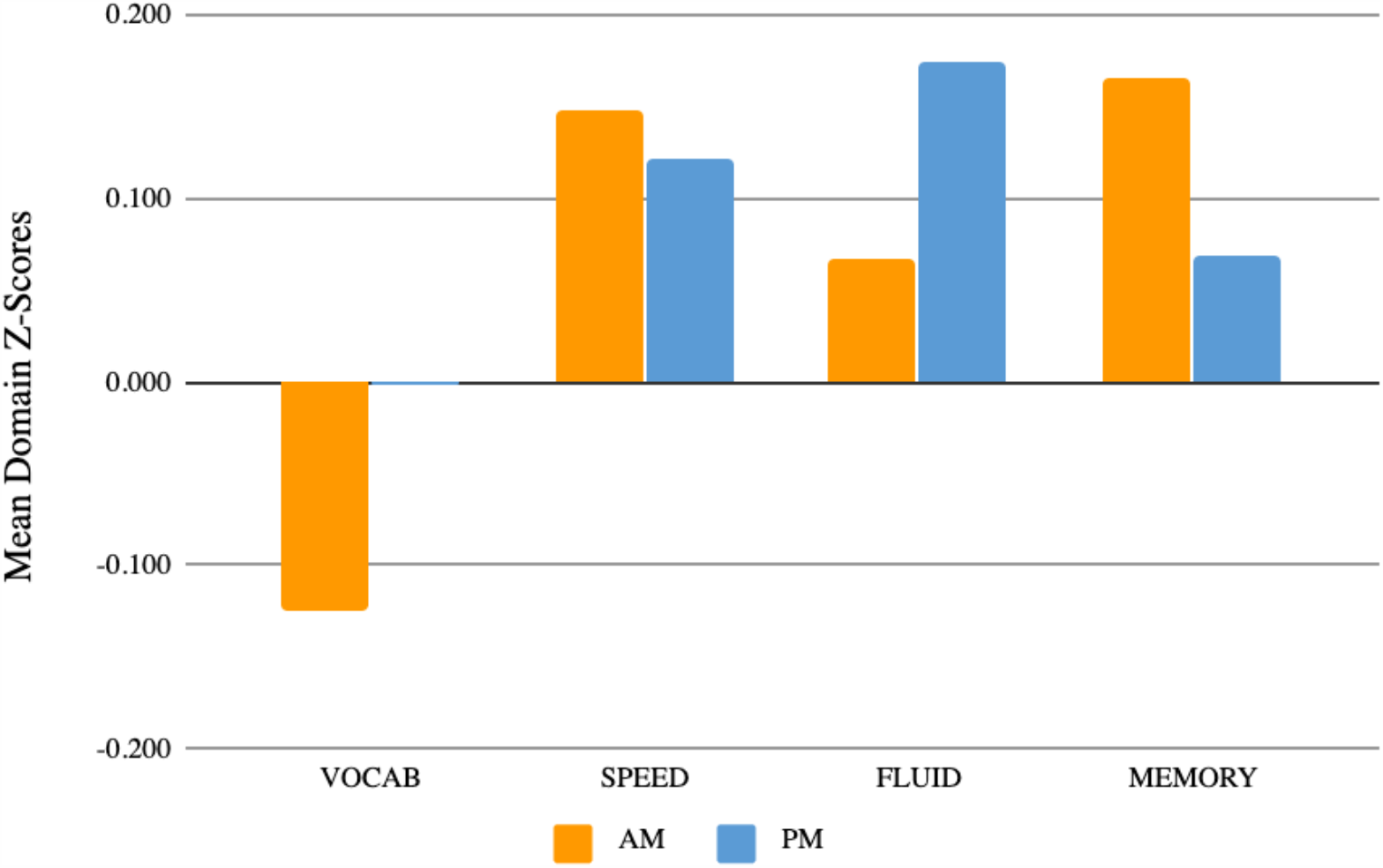
Model #1 interaction between Domain and time of day. For testing performed in either the morning (AM, orange) or afternoon (PM, blue), we present the mean Z score for each of the four tested cognitive domains (Vocabulary, Speed, Fluid Reasoning, and Memory).

**Figure 3:**
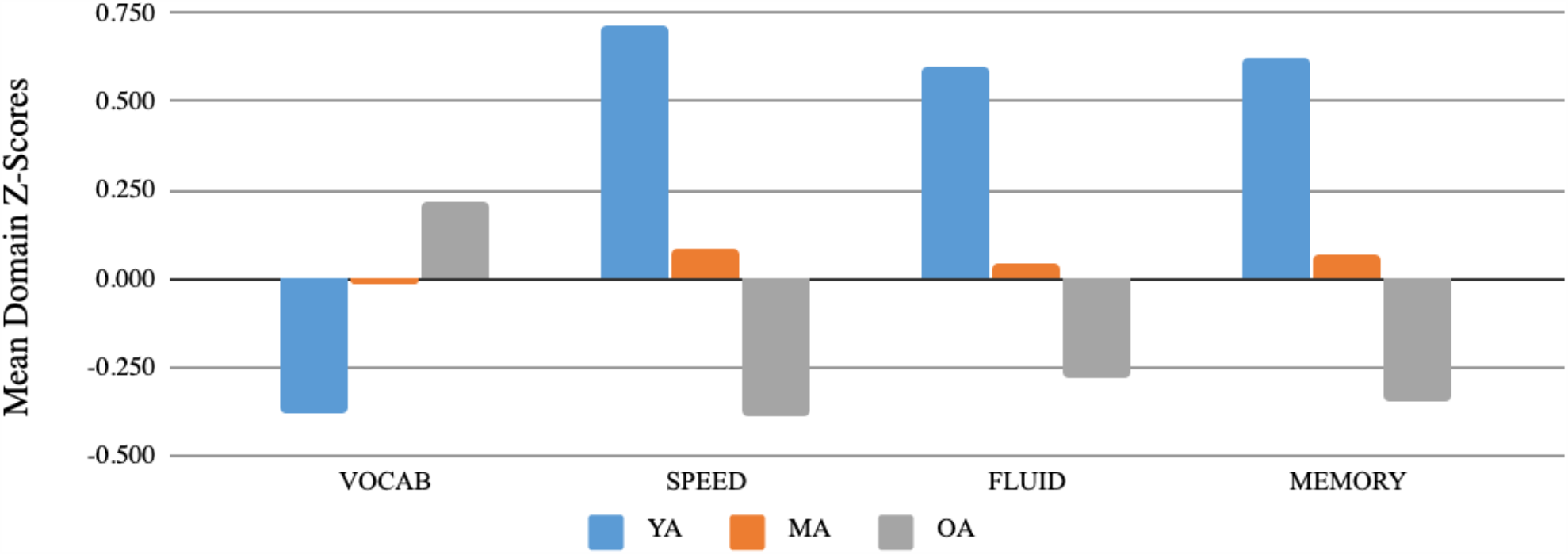
Model #1 interaction between Domain and Age. For each subgroup of subjects, we present the mean Z score for each cognitive domain (Vocabulary, Speed, Fluid Reasoning, and Memory). The Young Adults (YA) are shown in blue, the Middle Aged Adults (MA) in Orange and the Old Adults (OA) in gray.

To assess whether the analysis was limited by the way in which we defined the age groups, we performed a second analysis (Model #2) using a different binning strategy for participant age, classifying participants as either younger or older adults (**Figure 1b**). Specifically, a 4 (Domain: Vocabulary, Processing Speed, Fluid Reasoning, Episodic Memory) x 2 (Age Group: 20-49 years old, 50-80 years old) x 2 (AM: morning vs. afternoon testing) MANCOVA (covariates: years of education, gender) was utilized to probe these effects. It revealed significant main effects of Domain (F_3,1551_=31.989, p<.001) and Age Group (F_1,517_=106.507, p<.001), as well as a significant interaction between Domain and Age Group (F_3,1551_=116.603, p<.001). The main effect of time of day (F_1,517_=0.253, p=0.615), the interaction between Age Group and time of day (F_1,517_=0.226, p=0.635), the interaction between Domain and time of day (F_3,1551_=2.526, p=0.056), and the 3-way interaction among Domain, Age Group, and time of day (F_3,1551_=2.458, p=0.061) were not significant, as seen in Table 5 and Table 6. The main effect of Domain was identical to that observed in Model #1. The main effect of Age Group revealed that overall, adults under age 50 performed better on all tasks than adults age 50 plus (mean difference=0.525, p<.001). The interaction between Domain and Age Group showed that adults over age 50 performed better on Vocab tasks than adults under age 50 (t_526_=-5.886, p<.001), but for all other tasks, adults under age 50 performed better than adults over age 50 (Speed: t_526_=12.635, p<.001; Fluid: t_527_=10.398, p<.001; Memory: t_523_=10.937, p<.001). These results parallel those of model #1: there is no effect of time of day, and there is no effect of the interaction between time of day and age group on cognitive performance.

## Discussion

Results from the present study suggest that time of day was not related to cognitive performance, as assessed by tests measuring four domains: vocabulary, processing speed, fluid reasoning, and episodic memory. Within a sample of healthy adults, there was not a significant difference in participants’ performance on neuropsychological assessments based on whether they completed the tests in the morning or afternoon. In addition, since previous studies reported differences in synchrony with advancing age, we examined the effect of participant age in our analyses but found no evidence that the effect of diurnal rhythms on cognitive performance differed as a function of age. Nonetheless, one effect that showed a trend towards significance could warrant further investigation: Model #2 was the model closest to meeting our prespecified threshold of significance for an interaction among age, time of day, and task domain (p=0.056). This interaction trend showed that older adults may perform better on Vocabulary tasks but worse on Memory tasks in the afternoon, while younger adults may perform better on Fluid tasks in the afternoon (Figure 4). Although this was not significant, it may represent trends that could emerge as significant in a larger, or different, sample of subjects.

**Figure 4:**
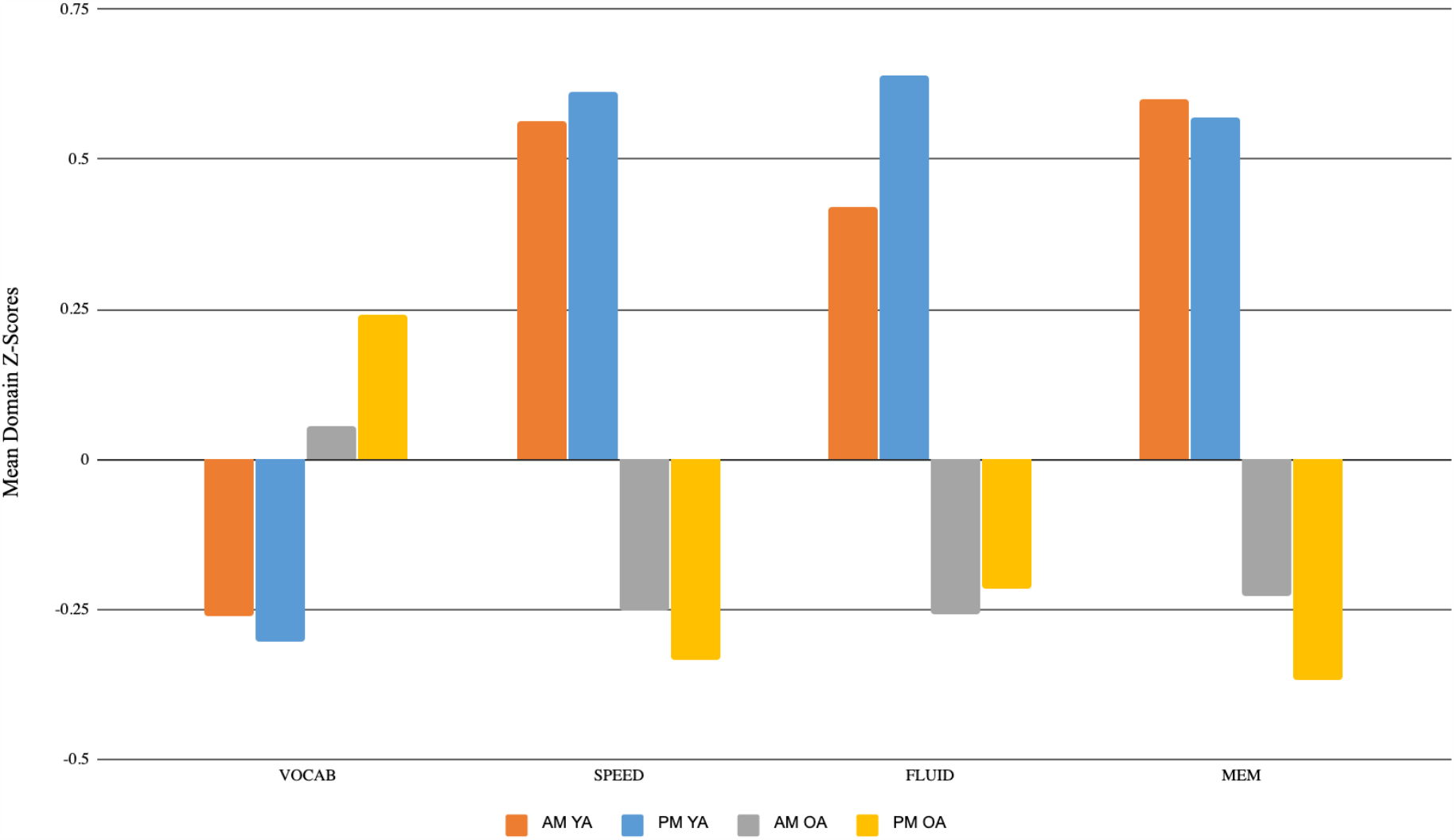
Model #2 three-way interaction between Domain, Age Group, and AM. For each subgroup of subjects, we present the mean Z score for each cognitive domain (Vocabulary, Speed, Fluid Reasoning, and Memory). The Young Adults (YA) are shown in blue, the Middle Aged Adults (MA) in Orange and the Old Adults (OA) in gray. There is no significant effect of time of day on performance for any of the four tested cognitive domains.

The results of this study are based on a large sample size and a diverse cognitive battery and thus make a significant contribution to the literature. Previous studies that evaluated the relationship between time of day, age, and performance on cognitive tasks returned inconsistent results. Most of these other studies were characterized by small sample sizes and single cognitive assessments, and thus their results may not be robust and are difficult to generalize. Some studies reported evidence that circadian effects may also vary with age: with older adults’ peak time of functioning being in the morning while younger adults’ peak time of functioning may tend to be later in the day (Adan & Almirall, 1990; Bodenhausen, 1990; Colquhoun, 1971; Folkard, Knauth, Monk, & Rutenfranz, 1976; Folkard, Weaver, & Wildgruber, 1983; Mecacci, Zani, Rocchetti, & Lucioli,1986; Wilson, 1990). Moreover, it was also noted that the patterns of performance throughout the day were still quite different for younger and older adults (May, Hasher, & Stoltzfus,1993). Finally, other investigators have also reported that they had not observed an effect of time of day on working memory performance (Sunram-Lea, 2001). Thus, we have made a contribution to the field by carefully examining the question of biological rhythms in a well-powered dataset using statistically rigorous methodology and four key cognitive domains. Our findings suggest that diurnal effects over the course of the work day are unlikely to have a meaningful effect on neuropsychological testing performance. If such effects exist, they would likely be very small since we could not detect them in 543 well-characterized subjects, or they may affect brain or cognitive functions not represented in these analyses. As far as the battery of standard tests that we examined, we can conclude that future studies may not need to adjust for time of day effects when utilizing these same neuropsychological tasks. Nonetheless, recording the time of testing in future studies would be useful to perform secondary analyses testing the findings of the present study, and could be examined along with our results in larger meta-analyses that might discover very small or specific effects on certain cognitive measures.

This study has several limitations. The number of old and young adults was not equal, and subjects came in whenever was most convenient for them (nonrandom assignment). This raises the possibility that participants selected the time of day that was more advantageous for their performance. That being said, based on these findings, for those studies that allow participants to select the timing of their appointment, time of testing may not be necessary to account for in analyses comparing younger and older adults’ performance. The distribution of people having AM or PM appointments was also similar but not equal. Further, there are other important variables that were not recorded and could be important, such as each participant’s circadian clock, or whether they are a “morning” or “evening” person. This is important because it was previously noted that patterns of performances throughout the day are different for younger and older adults (May, Hasher, & Stoltzfus,1993). Finally, we did not record the length of the appointment, so we do not know whether the length of the appointment has any effect on the subject’s performance of different cognitive tasks.

A more targeted study that is optimally designed to test a time of day effect would optimally include an equal number of older and younger adults tested in the AM or PM, and would randomize time assignments. It should also include a Morningness-eveningness questionnaire. Finally, to generate a more comprehensive study of domain-specific effects, one could evaluate scores on specific cognitive assessments and how they correlate with age and time of day rather than solely examining domain-based z-scores from neuropsychological tests because it is possible that only selected individual tasks may be sensitive to diurnal rhythms. Further, future studies could also monitor one’s sleep schedule so that we can fully understand each participant’s individual sleep cycle and circadian clock, and see how these more detailed measurements affect cognitive performance.

Overall, the analyses reported here suggest that time of day does not meaningfully influence performance on the tasks administered in the present study. More generally, our observation is reassuring that time of day does not have to be considered as a key variable when designing new studies; given that human studies include large samples of participants who lead busy lives, having to control for the time of day in scheduling participants could create significant hardship in executing a project, particularly if the time of day had to be personalized because of age or other factors. With these results in hand, we can now further explore the role of these important biological rhythms on cognition by exploring this question in different ways, such as testing performance in other tasks that we did not test, and whether more extreme differences in time - such as morning vs nighttime - uncover differences in cognitive function.

## Acknowledgements

We thank the participants in the RANN and CR studies for their time in contributing to these studies. This research was supported by two grants from the studies National Institutes on Aging (RANN study: RF1AG038465, principal investigator: YS; Cognitive Reserve study: RO1AG026158, principal investigator: YS). There are no conflicts of interest.

